# Extradomain-B Fibronectin is a molecular marker of invasive breast cancer cells

**DOI:** 10.1101/743500

**Authors:** Amita M. Vaidya, Helen Wang, Victoria Qian, Zheng-Rong Lu

## Abstract

Breast tumor heterogeneity is a major impediment to oncotherapy. Tumor cells undergo rapid clonal evolution, thereby acquiring significant growth and invasive advantages. The absence of specific markers of these high-risk tumors precludes efficient therapeutic and diagnostic management of breast cancer. Given the critical function of tumor microenvironment in the oncogenic circuitry, we sought to determine the role of the extracellular matrix oncoprotein, extradomain-B fibronectin (EDB-FN), as a molecular marker of aggressive cancers. High-risk invasive cell lines generated from relatively less invasive MCF7 and MDA-MB-468 breast cancer cells by long-term TGF-β treatment and chemoresistance demonstrated hybrid epithelial-mesenchymal phenotype, enhanced motility, and significantly elevated EDB-FN levels in 2D- and 3D-cultures. To determine if EDB-FN could serve as a therapy-predictive marker, the invasive cell lines were treated with MK2206-HCl, a pan-AKT inhibitor. Phospho-AKT depletion reduced EMT and invasion of the populations, with a concomitant decrease in EDB-FN expression, partly through the phosphoAKT-SRp55 pathway, demonstrating that EDB-FN expression is strongly associated with high-risk breast cancer. EDB-FN is a promising molecular marker for accurate detection, differential diagnosis, and non-invasive therapeutic surveillance of aggressive breast cancer.

**Summary Statement:** Dynamic changes in invasive properties of breast cancer cells directly influence extradomain-B fibronectin levels, suggesting its potential role as a molecular marker for active surveillance and therapeutic monitoring of breast cancer.

## Introduction

Breast cancer (BCa) is a devastating disease that accounts for 41,000 deaths each year in the US (Siegel et al., 2018). Although the survival rate for patients with localized BCa is close to 99%, it declines precipitously in patients with distant metastases and drug resistance (Siegel et al., 2018, DeSantis et al., 2017). A major stumbling block in the clinical management of the disease is tumor heterogeneity, which plays a role in the dynamic nature of BCa progression (Baird and Caldas, 2013). Whole genome sequencing and profiling studies have demonstrated that breast tumors of the same histological subtype exhibit distinct molecular portraits and discrete trajectories in individual BCa patients at different stages (Tsang and Tse, 2019, Eliyatkin et al., 2015, Baird and Caldas, 2013). Stochastic mutations, genome instability, and clonal evolution arising from selective pressures from genetic, epigenetic, environmental, and therapeutic stimuli result in the emergence of high-risk tumor populations with significant growth and invasive advantages (Eccles et al., 2013, Baird and Caldas, 2013). This extensive spatial and temporal diversity within primary and metastasized tumors directly influences diagnostic, therapeutic, and prognostic outcomes (Eccles et al., 2013). In the absence of markers specific to the metastatic and invasive properties of tumors, current imaging modalities including MRI, PET, and CT are limited in their ability to detect and differentiate between low-risk and high-risk tumors (Bleyer and Welch, 2012). These facts underscore the need for the discovery and characterization of suitable molecular markers that can facilitate non-invasive detection, risk-stratification, active surveillance of breast neoplasms, and timely assessment of therapeutic response, despite their dynamic nature.

The tumor extracellular matrix (ECM) plays a critical role in all aspects of tumor progression, by relaying oncogenic signals between the tumor cells and the tumor microenvironment (TME) and by supporting growth, apoptotic escape, migration, inflammation, and immune evasion (Balkwill et al., 2012). Fibronectin (FN1), an integral component of normal and tumor ECM, is an essential glycoprotein that regulates adhesion, motility, growth and development (Pankov and Yamada, 2002). Its alternative splice variant called extradomain-B fibronectin (EDB-FN), however, is known to be expressed during malignant transformation, and is generally absent from healthy adult tissues (White et al., 2008, Han and Lu, 2017). Multiple lines of evidence show that EDB-FN is associated with epithelial-to-mesenchymal transition (EMT), cancer cell stemness, proliferation, angiogenesis, and metastasis, all of which reflect tumor aggressiveness (Petrini et al., 2017, Sun et al., 2015, Tavian et al., 1994, Coltrini et al., 2009, Ventura et al., 2018, Han et al., 2018). Clinical studies demonstrate the presence of EDB-FN in patients with lung, brain, colorectal, ovarian, and thyroid cancers (Santimaria et al., 2003, Menzin et al., 1998, Giannini et al., 2003). The overexpression of EDB-FN is also correlated with histological grade in mammary tumors (Loridon-Rosa et al., 1990) and with poor survival in oral carcinoma patients (Lyons et al., 2001), suggesting its potential role as a marker for multiple neoplasms.

An added layer of complication is that even among the same cancer type, EDB-FN expression profiles are distinct and specific to the molecular and functional characteristics of the cells or tissues. For example, using an EDB-FN-specific peptide probe, ZD2-Cy5.5, we previously showed that invasive cancer cell lines, e.g., PC3 (prostate) and MDA-MB-231 (hormone receptor-negative breast cancer), are EDB-FN-rich, while the less invasive cancer cell lines, e.g., LNCaP (prostate) and MCF7 (hormone receptor-positive breast) exhibit significantly lower EDB-FN levels (Han et al., 2015, Han et al., 2017a, Han et al., 2018, Han et al., 2017b). This differential expression of EDB-FN was exploited for differentially diagnosing invasive prostate and breast cancer tumors from the non-invasive xenografts using EDB-FN-targeted MRI contrast agents (Ayat et al., 2018, Han et al., 2017a, Han et al., 2017b). Other independent groups have also used EDB-FN as a molecular target for targeted imaging and therapeutic delivery for various types of cancer (Sun et al., 2014, Ye et al., 2017, Han et al., 2019).

Given the high degree of tumor plasticity, it is evident that different selective pressures, environmental and experimental stimuli will bring about distinct changes in the TME and EDB-FN expression, which would in turn influence the clinical outcomes of EDB-FN-targeted imaging and therapeutic interventions. Here, we sought to determine the changes in EDB-FN expression patterns following application of two different selective pressures on non-invasive, low EDB-FN-expressing breast cancer cells and their consequent evolution into invasive high-risk populations. To this end, significant survival advantage was conferred on two breast cancer cell lines, MCF7 and MDA-MB-468, by treating them with the cytokine TGF-β and chemotherapeutic drugs to induce stochastic alternations and clonal evolution. The resulting populations were also treated with a highly specific AKT inhibitor to further assess for changes in EDB-FN levels in correlation to the response of the high-risk cells to targeted therapy.

## Results

### EDB-FN expression is significantly elevated in breast cancer

As a critical ECM component, FN1 is overexpressed in multiple cancer types (Bae et al., 2013, Suer et al., 1996, Saito et al., 2008, Menendez et al., 2005). Here, the expression of the FN1 transcript containing the EDB-FN exon (ENST00000323926) was assayed by performing differential gene expression analysis on RNA-Seq patient data from TCGA database. As shown in **Fig. 1A**, breast tumors demonstrated significant overexpression of the EDB-FN transcript, compared to normal breast tissues. Next, the endogenous levels of EDB-FN mRNA were determined across a panel of cell lines representing the multiple molecular subtypes of breast cancer (Holliday and Speirs, 2011, Arnedos et al., 2012). As shown in **Fig. 1B**, the least invasive hormone receptor-positive (HR^+^) MCF7 cells showed the lowest expression of EDB-FN. The more invasive triple-negative (HR^-^) breast cancer lines showed significant upregulation of EDB-FN levels, with 4-fold increase in MDA-MB-468 cells, 8-10-fold increase in BT549 and MDA-MB-231 cells, and a 555-fold increase in Hs578T cells. These results demonstrate that the oncofetal EDB-FN isoform is highly expressed in malignant breast phenotypes.

**Figure 1.**
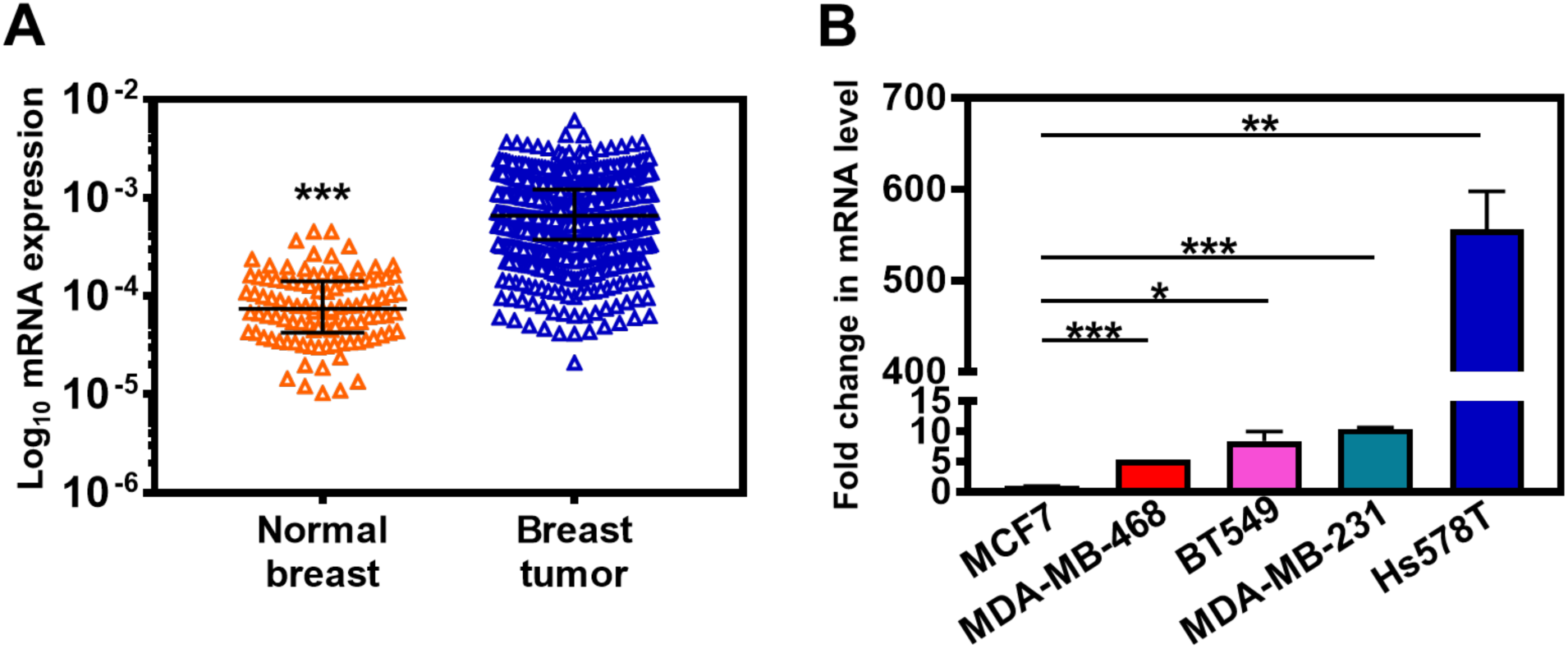
EDB-FN overexpression in breast cancer. **A.** Differential gene expression analysis performed on patient data from the TCGA database shows significant overexpression of EDB-FN transcript (ENST00000323926) in breast tumor samples (n=790) compared to normal breast tissue samples (n=104). Scatter dot plot denotes FPKM values and median with interquartile range, *p<0.0001 using unpaired 2-tailed *t*-test. **B.** Endogenous expression of EDB-FN in different subtypes of breast cancer cell lines as measured by qRT-PCR. Triple negative MDA-MB-468, BT549, MDA-MB-213, and Hs578T cells exhibit significantly elevated levels of EDB-FN mRNA, compared to the hormone receptor-positive MCF7 cells. 18S expression was used as standard. Bars denote mean ± sem (n=3). Unpaired 2-tailed *t*-test used to determine p values (*p<0.05, **p<0.01, ***p<0.001).

### Growth and morphology changes in 2D- and 3D-cultured breast cancer cells with TGF-β treatment and drug resistance

To assess the changes in EDB-FN expression levels when breast cancer cells gain significant survival advantages, the two cell lines with the lowest EDB-FN expression and epithelial phenotype, namely MCF7 and MDA-MB-468 cells, were chosen. Two selective pressures were applied: 1) long-term treatment with TGF-β (5 ng/mL) to induce EMT (Xu et al., 2009) to generate MCF7-TGFβ and MDA-MB-468-TGFβ cells and 2) acquired chemoresistance to Palbociclib, a cyclin-dependent kinase (CDK) inhibitor (Chen et al., 2018), and to Paclitaxel, an anti-microtubule agent (Barbuti and Chen, 2015), to generate MCF7-DR and MDA-MB-468-DR cells, respectively. The parent and derivative cell lines were characterized for their morphology, and molecular and functional phenotypes.

As shown in **Fig 2A**, MCF7 cells demonstrate a typical epithelial morphology in 2D culture. Long-term treatment with TGF-β and development of resistance to Palbociclib resulted in morphological changes to a more mesenchymal phenotype, which was more pronounced in the MCF7-DR cells than in MCF7-TGFβ cells. On the other hand, the MDA-MB-468 cells did not exhibit overt changes in morphology with TGF-β treatment and development of resistance to Paclitaxel. However, the MDA-MB-468-TGFβ showed increased growth rate compared to the parent MDA-MB-468 cells.

**Figure 2.**
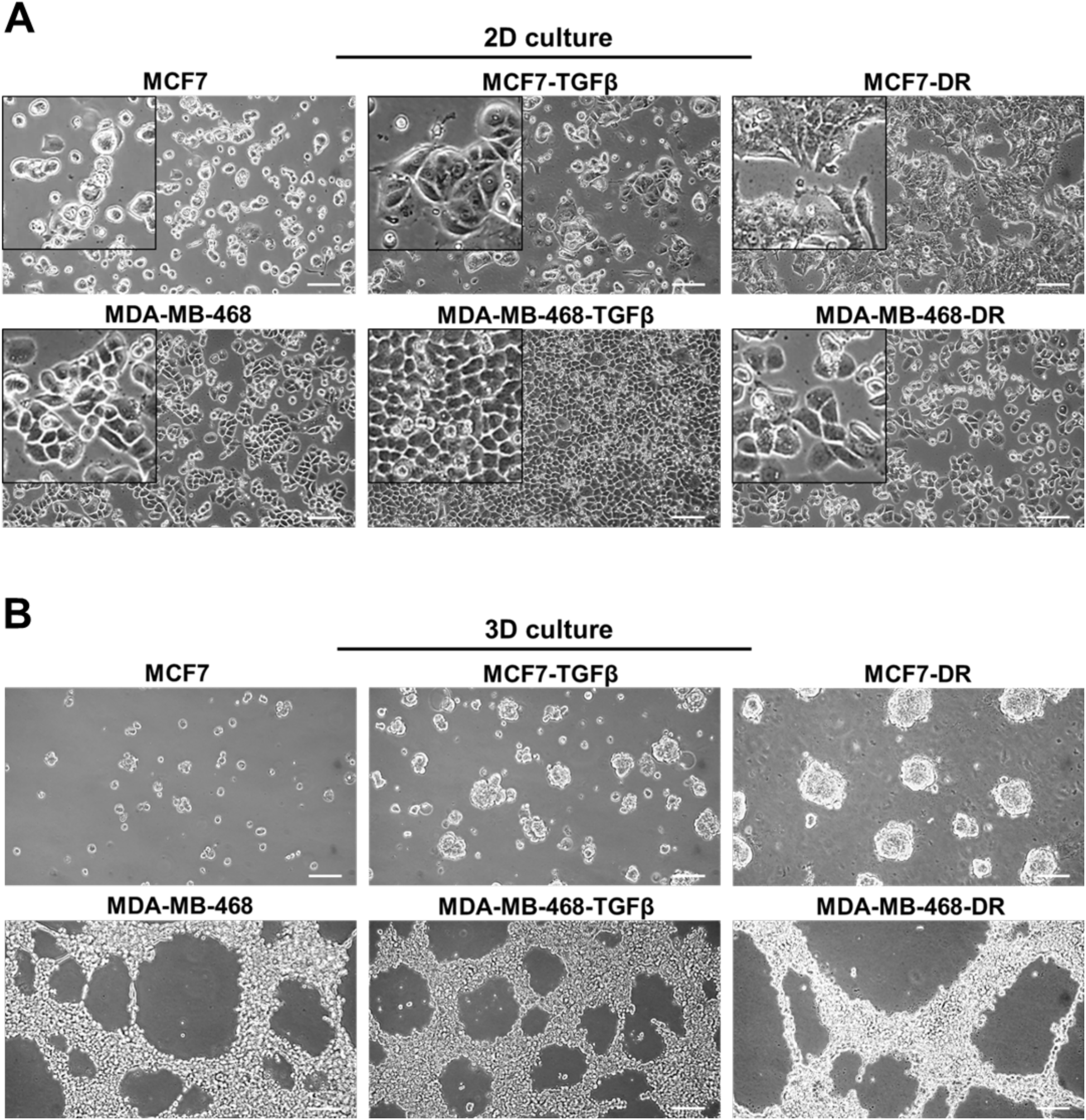
Growth and morphology changes in breast cancer cells in 2D and 3D culture. MCF7 and MDA-MB-468 cells were cultured in 5 ng/mL TGF-β for 7-15 days to obtain MCF7-TGFβ and MDA-MB-468-TGFβ, cells respectively. MCF7-DR and MDA-MB-468-DR cells were obtained by inducing resistance to 500 nM Palbociclib and 100 nM Paclitaxel respectively. **A.** MCF7-TGFβ and MCF7-DR cells show distinct morphological changes, with a more mesenchymal phenotype, in 2D culture and increased ability to form spheroids in 3D Matrigel culture, compared to their parent lines. **B.** MDA-MB-468-TGFβ and MDA-MB-468-DR cells do not show visible morphological changes in 2D culture and form similar proliferative networks in 3D Matrigel culture, compared to the parent counterparts.

In addition to 2D culture, the cells were grown in Matrigel to facilitate the establishment of a conducive ECM. As shown in **Fig. 2B**, in 3D culture, the low-risk HR^+^ MCF7 cells showed negligible tumor spheroid formation while the more invasive MDA-MB-468 cells showed proliferative network formation. The MCF7-TGFβ and MCF7-DR cells formed tumor spheroids, unlike the parent MCF7 cells, while the MDA-MB-468-TGFβ and MDA-MB-468-DR cells formed similar proliferative networks as their parent counterparts. These results highlight the different properties of each cancer cell type and its distinct response to external mitogenic stimuli.

### Increased migration in breast cancer cells with TGF-β treatment and drug resistance

Next, we analyzed the molecular and functional changes in the TGF-β-treated and drug-resistant breast cancer cells. TGF-β is a potent inducer of EMT, a critical step towards initiation of metastasis (Fedele et al., 2017). Similarly, the signaling programs of EMT and drug resistance are intricately related, where EMT-like molecular signature can antagonize chemotherapy in breast cancer (Huang et al., 2015). Consequently, the mRNA and protein expression of the common EMT markers, N-cadherin (N-cad), E-cadherin (E-cad), and Slug (invasion marker), was tested in the derivative cell lines. As shown in **Figs. 3A-C**, MCF7-TGFβ and MCF7-DR cells showed increased EMT, evidenced by reduced E-cad and increased N-cad and Slug mRNA levels, in comparison to MCF7 cells. MDA-MB-468-TGFβ and MDA-MB-468-DR cells, however, did not demonstrate significant changes in E-cad and N-cad mRNA levels, compared to their parent cells. The mRNA expression of Slug increased only in the TGF-β-treated MDA-MB-468 cells but not in MDA-MB-468-DR cells (**Fig. 3C**).

**Figure 3.**
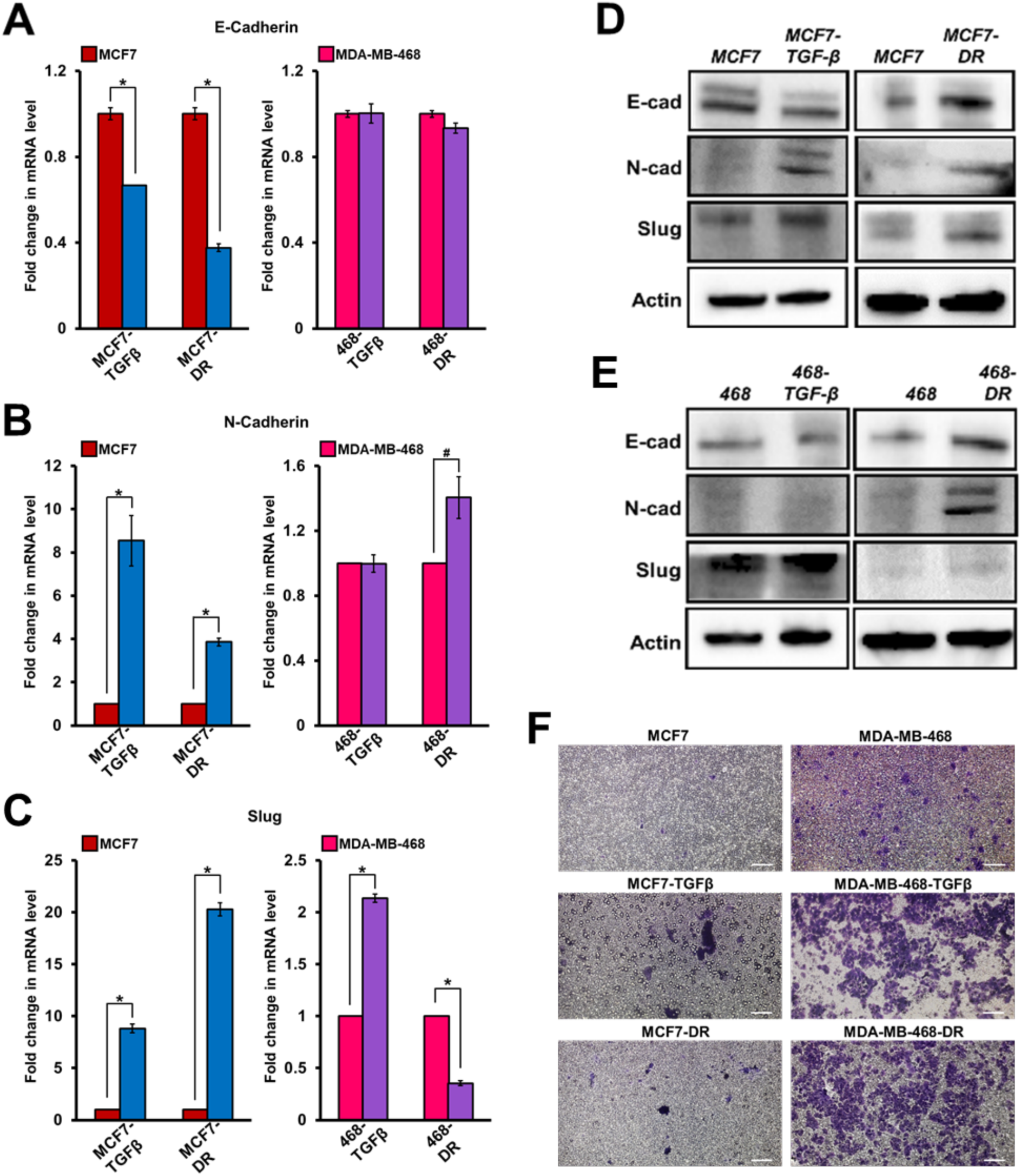
Increased E-M phenotype and enhanced migration in breast cancer cells with TGF-β treatment and drug resistance. qRT-PCR analysis demonstrates mRNA expression of **A.** E-cadherin, **B.** N-cadherin, and **C.** Slug in MCF7 and MDA-MB-468 following TGF-β treatment and development of drug resistance. Western blot analysis for EMT markers shows protein expression of E-cad, N-cad, and Slug in **D.** MCF7-TGFβ and MCF7-DR and **E.** MDA-MB-468-TGFβ and MDA-MB-468-DR cells, compared to the parent MCF7 and MDA-MB-468 cells, respectively. **F.** Transwell assay shows higher invasive potential of TGF-β-treated and drug-resistant MCF7 and MDA-MB-468 cells, evidenced by the increase in number of crystal violet-stained migrated cells. Bars denote mean ± sem (n=3). Unpaired 2-tailed *t*-test shows *p<0.05, #p=0.06. Scale bar = 100 µm.

At the protein level, both MCF-TGFβ and MCF7-DR cells showed upregulated N-cad and Slug expression while the E-cad expression did not significantly decrease (**Fig. 3D**), indicating that the MCF7 cells gain a partial EMT-like phenotype with TGF-β treatment and development of drug resistance. The MDA-MB-468-TGFβ cells showed no changes in E-cad and N-cad and a moderate increase in Slug levels while the MDA-MB-468-DR cells showed increased N-cad and Slug expression, compared to the parent cells (**Fig. 3E**). These results indicate that while both the MDA-MB-468-TGFβ and MDA-MB-468-DR cells overexpress the migratory protein Slug, the former did not undergo EMT with TGF-β treatment while the latter possibly gained a hybrid E-M phenotype with drug resistance.

The treated cell populations were analyzed for their ability to invade through a layer of matrigel coated in transwell inserts. As shown in **Fig. 3F**, both the TGF-β treatment and drug resistance conferred significant invasive advantage on the MCF7 and MDA-MB-468 cells, rendering them more motile than their parent counterparts, as seen by the increased number of crystal violet-stained migrated cells. Indeed, recent studies have revealed that the partial or hybrid E-M phenotype is attributed to the tumor cell plasticity and is extremely favorable for metastatic dissemination (Kroger et al., 2019, Saitoh, 2018).

### Increased EDB-FN expression in breast cancer cells with TGF-β treatment and drug resistance

The potential role of EDB-FN as a molecular marker for breast cancer aggressiveness was then determined in TGF-β-treated and drug-resistant MCF7 and MDA-MB-468 cells. The expression of EDB-FN in 3D-cultured cells was analyzed using fluorescent-labeled EDB-FN-specific peptide ZD2-Cy5.5 (Han et al., 2015). As shown in **Fig. 4A**, endogenous EDB-FN expression in MDA-MB-468 cells is higher than that in the MCF7 cells, consistent with the mRNA levels in **Fig. 1B**, and their invasive ability in **Fig. 3F**. Treatment with TGF-β and acquired drug resistance led to a significant increase in EDB-FN expression in the 3D-cultured MCF7 and MDA-MB-468 cells. The EDB-FN-specific binding of the ZD2-Cy5.5 probe was confirmed by EDB-FN knockdown experiments in MDA-MB-468-DR cells, where ECO/siEDB nanoparticle treatment abrogated the ZD2-Cy5.5 binding (**Fig. 4B**). The peptide binding results from 3D culture were also corroborated by qRT-PCR analysis, which showed over 3-fold and 10-fold increase in EDB-FN expression in MCF7-TGFβ and MCF7-DR cells and about 4-fold increase in MDA-MB-468-TGFβ and MDA-MB-468-DR cells, over their respective counterparts (**Fig. 4C**). These results indicate that the acquisition of partial EMT and invasive properties by breast cancer cells results in upregulation of EDB-FN. Thus, EDB-FN overexpression is strongly upregulated in invasive breast cancer cells and in the breast cancer cells that evolve from low-risk ones.

**Figure 4.**
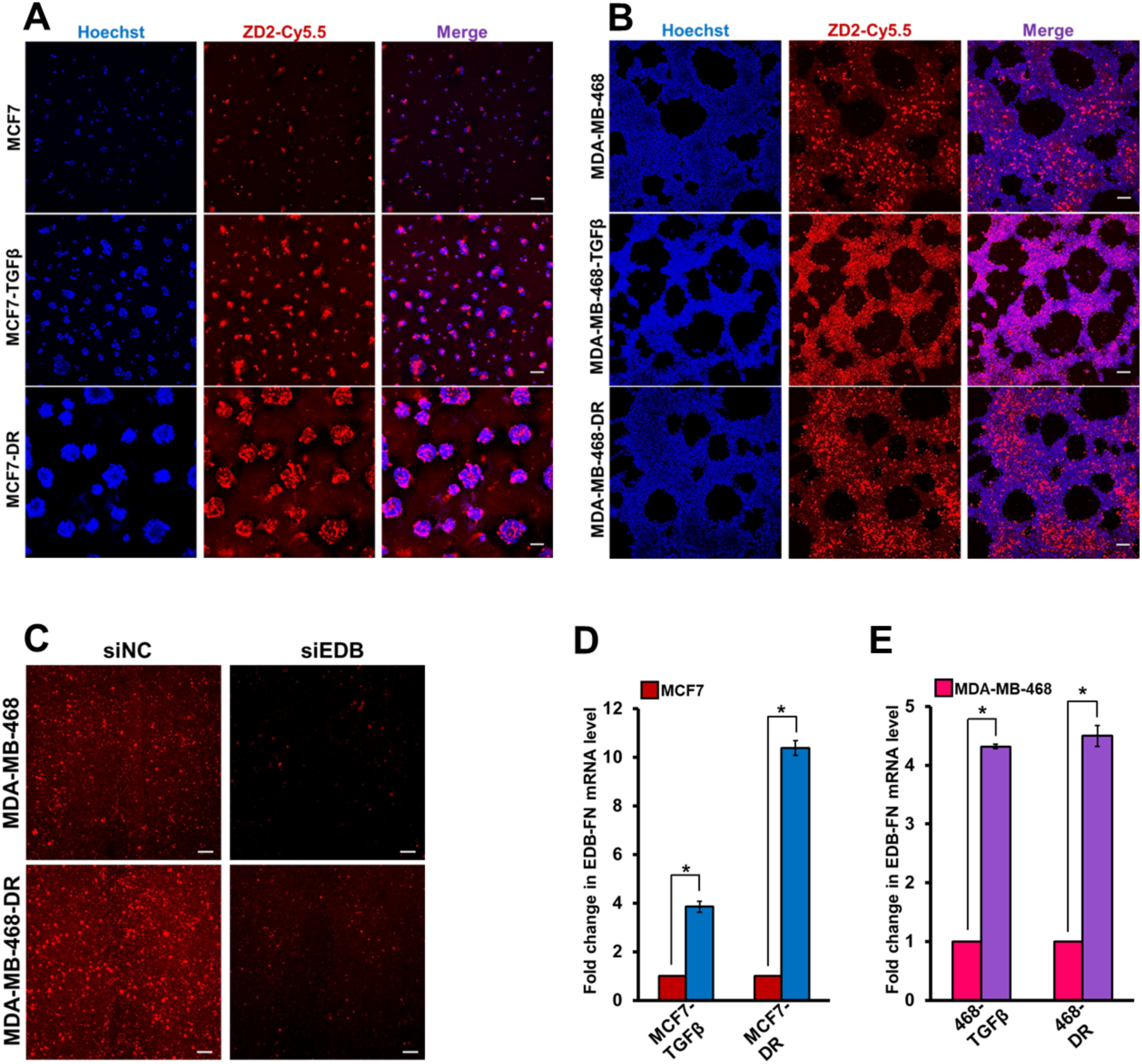
Increased EDB-FN expression in breast cancer cells with TGF-β treatment and drug resistance. **A.** ZD2-Cy5.5 staining of 3D cultures of breast cancer cells shows significantly increased EDB-FN expression in **A.** MCF7-TGFβ and MCF7-DR and **B.** MDA-MB-468-TGFβ and MDA-MB-468-DR cells, compared to the parent MCF7 and MDA-MB-468 cells, respectively. **C.** ZD2-Cy5.5 binding to EDB-FN is abolished with RNAi of EDB-FN using ECO/siEDB-FN nanoparticles, confirming EDB-FN-specific binding of ZD2-Cy5.5. ECO/siLuc nanoparticles used as negative control. qRT-PCR analysis shows significantly upregulated mRNA expression of EDB-FN in **D.** MCF7-TGFβ and MCF7-DR and **E.** MDA-MB-468-TGFβ and MDA-MB-468-DR cells, compared to the parent MCF7 and MDA-MB-468 cells, respectively (*p<0.05). Bars denote mean ± sem (n=3). Unpaired 2-tailed *t*-test shows *p<0.05. Scale bar = 100 µm.

### Therapeutic ablation of AKT in invasive breast cancer cells decreases invasion and EDB-FN expression

Non-invasive therapeutic monitoring of tumor response to oncostatic drugs is crucial to facilitate decision making and timely interventions (Eccles et al., 2013). To test whether EDB-FN is a therapy-predictive marker and if its expression correlates with changes in the invasive potential of breast cancer cells, the TGF-β-treated and drug-resistant MCF7 and MDA-MB-468 cells were treated with MK2206-HCl, a highly specific pan-AKT inhibitor proven to suppress PI3K/AKT signaling-induced tumor cell proliferation (Hirai et al., 2010). The PI3K/AKT signaling is a major signal transduction cascade implicated in the progression, metastasis, and drug resistance of multiple cancers (Agarwal et al., 2014).

The upregulation of the mitogenic AKT signaling axis in the aggressive TGF-β-treated and drug-resistant MCF7 and MDA-MB-468 cell populations was first confirmed by testing for the levels of phosphorylated AKT (T308 and S473) and total AKT (**Fig. 5A**). Both the phospho-AKT-T308 and phospho-AKT-S473 levels were strongly upregulated in the TGF-β-treated and drugresistant MCF7 and MDA-MB-468 cells, in addition to increased expression of total AKT in the MDA-MB-468-TGFβ cells. Treatment of the invasive cell derivatives with MK2206-HCl resulted in robust inhibition of phospho-AKT (T308 and S473), as shown in **Fig. 5B**, and subsequently reduced their invasive potential (**Fig. 5C-D**). This was accompanied by a significant decrease in the expression of EDB-FN in the EDB-FN-overexpressing TGF-β-treated and drug-resistant MCF7 and MDA-MB-468 cells, demonstrated by the significant decrease in the mRNA levels (**Fig. 5E-F**) and reduced ZD2-Cy5.5 staining in 3D cultures (**Fig. 5G-H**), compared to the DMSO-treated controls. Additionally, the invasive cells also decreased N-cad (prominent in MDA-MB-468 cells) and Slug expression (**Fig. 5I**), in consistence with the diminished EDB-FN expression and invasiveness. These results demonstrate a positive association between EDB-FN and aggressiveness of breast cancer cell lines, and potential correlation of altered EDB-FN expression with response to therapeutic interventions.

**Figure 5.**
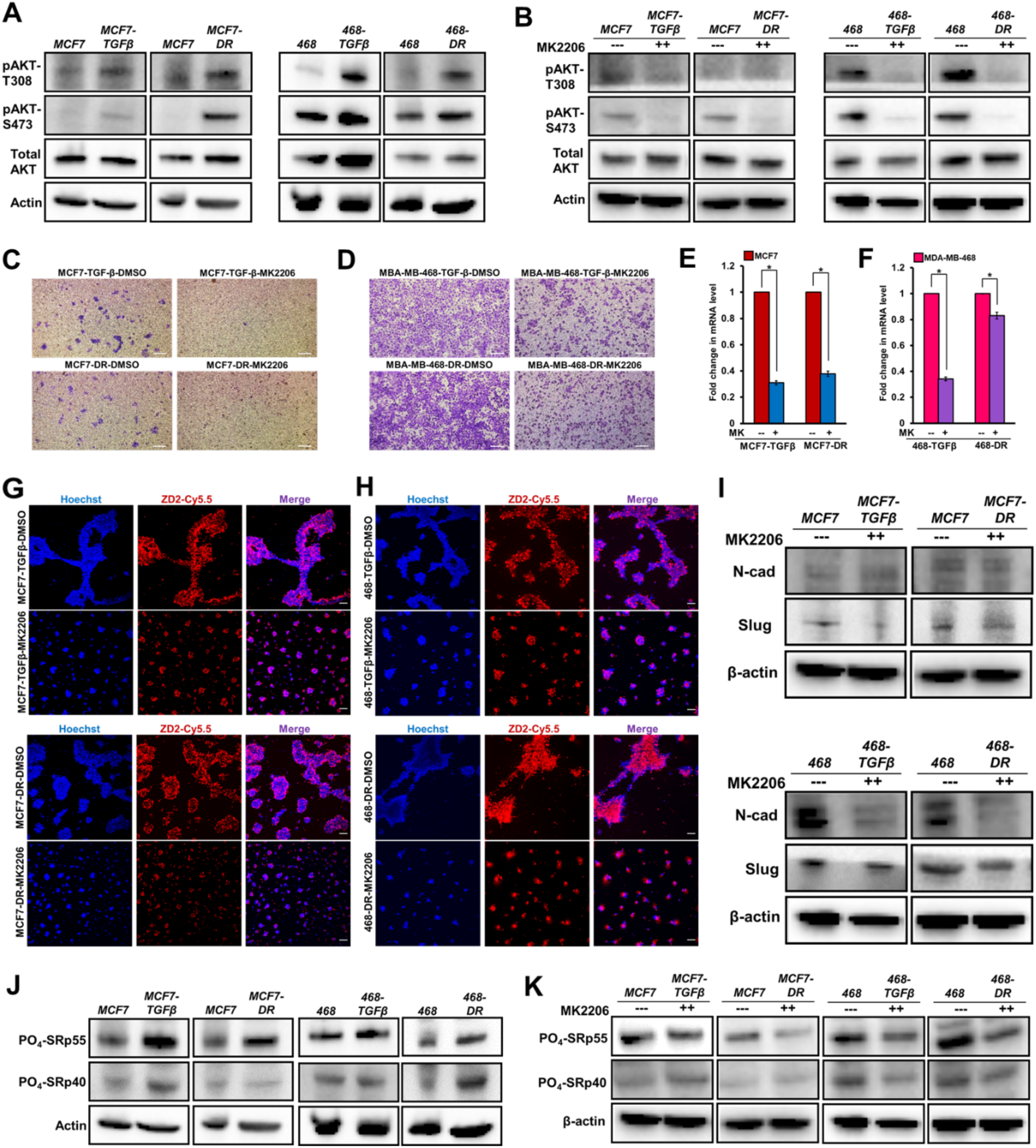
Therapeutic ablation of AKT in invasive TGF-β-treated and drug-resistant breast cancer cells reduces their invasion and EDB-FN overexpression. **A.** TGF-β treatment and drug resistance upregulate phospho-AKT signaling in MCF7 and MDA-MB-468 cells. **B.** Treatment of cells with AKT inhibitor MK2206-HCl (2 uM for MCF7 and 4 uM for MDA-MB-468 cells) for 2 days results in inhibition of phospho-AKT signaling in the invasive breast cancer cell lines. Inhibition of phospho-AKT signaling by MK2206-HCl demonstrates **C and D**. reduced invasive potential of the invasive lines along with decreased expression of EDB-FN at **E and F.** mRNA levels and in **G and H.** 3D culture. **I.** Phospho-AKT depletion leads to reduced expression of N-cad and Slug in the invasive MCF7 and MDA-MB-468 breast cancer lines. **J.** TGF-β treatment and drug resistance upregulate phosphorylation of SRp55 in MCF7 and MDA-MB-468 cells. **K.** This upregulation is abolished with MK2206-HCl-mediated depletion of phospho-AKT signaling, suggesting a potential role of SRp55 in the inclusion of EDB-FN exon. Bars denote mean ± sem (n=3). Unpaired 2-tailed *t*-test shows *p<0.05. Scale bar = 100 µm.

Previous studies have implicated the role of phosphoAKT-SRp40 pathway in the alternative splicing-mediated regulation of EDB-FN expression (Bordeleau et al., 2015). To determine the mechanism of EDB-FN upregulation by TGF-β treatment and drug resistance, we examined the expression of phosphorylated SR proteins (SRp55 and SRp40) in the invasive breast cancer cells before and after treatment with MK2206-HCl. As shown in **Fig. 5J**, MCF7-TGFβ, MCF7-DR, MDA-MB-468-TGFβ, and MDA-MB-468-DR cells upregulate the expression of SRp55, while only MCF7-TGFβ and MDA-MB-468-DR upregulate SRp40, compared to their respective parent counterparts. The upregulation of SRp55 was abolished in the four cell lines with MK2206-HCl treatment (**Fig. 5K**), suggesting that EDB-FN upregulation in these invasive cells could be controlled, at least in part, through the phosphoAKT-SRp55 signaling pathway.

## Discussion

Numerous blood biomarkers including CA 15.3, carcinoembryonic antigen (CEA), CA125 and imaging modalities like ultrasound, mammography, MRI, PET, and CT are routinely used to detect primary breast tumor disease and recurrence and to assess therapeutic response (Khatcheressian et al., 2013, Bayo et al., 2018). However, they are limited in their ability to differentially diagnose and risk-stratify the disease, with high rates of false positive diagnoses (Othman et al., 2015), underscoring the need for specific markers to accurately detect highly invasive and metastatic breast tumors, and to distinguish them from low-risk indolent ones. Moreover, breast tumors frequently exhibit intrinsic or acquired resistance to chemotherapy and targeted drugs (Huang et al., 2015). In the absence of suitable molecular markers, active surveillance and monitoring of the efficacy of chemotherapeutic interventions and timely detection of the emergence of resistant phenotypes forms another obstacle to patient treatment.

To address these concerns, this study investigated the dynamic changes in the ECM oncoprotein EDB-FN in conjunction with the dynamic changes in the invasive potential of breast cancer cells. We found that invasive cells that evolve from low-risk cancer cells exhibit partial EMT-like phenotypes and overexpress EDB-FN; conversely, impeding the invasive abilities of these high-risk cancer cells with a targeted drug abolishes their EDB-FN overexpression, demonstrating a direct correlation between EDB-FN levels and the invasiveness of breast cancer cells. Moreover, to our knowledge, this is the first study to report that EDB-FN is upregulated with development of drug resistance in breast cancer cells.

Between the two different breast cancer lines used in this work, the endogenous EDB-FN level in the least aggressive HR^+^ MCF7 cells is significantly lower than that in the more aggressive HR^-^ MDA-MB-468 cells, despite both lines exhibiting an epithelial phenotype. Induction of drug resistance and long-term TGF-β treatment led to distinct changes in the molecular phenotypes of thetwo cell lines, possibly through distinct signaling mechanisms. Nevertheless, the emergent invasive populations became more aggressive than their parent cells, with a pre-metastatic hybrid E-M phenotype and increased phospho-AKT signaling. It is also likely that each of these cell lines acquired their survival advantages through additional here-to-fore unstudied mechanisms. Nevertheless, all of the cells with acquired invasiveness presented elevated EDB-FN expression irrespective of the signaling mechanisms. Conversely, when the aggressive cells were treated with targeted therapy, their invasive potential diminished with a concomitant reduction in EDB-FN expression, indicating the role of EDB-FN as a marker for active surveillance and monitoring of therapeutic efficacy.

The precise mechanism of EDB-FN upregulation in invasive cells remains an enigma. At the genetic level, EDB-FN is generated by alternative splicing event, resulting in the inclusion of the EDB exon in the FN1 transcript, a process controlled by SR (Ser- and Arg-rich) proteins of the splicing regulator family (White et al., 2008, Huh and Hynes, 1994). Since alternative splicing is indispensable for the formation of the EDB-FN isoform, the participation of the SR proteins in this process is inevitable. However, there is limited research on the underlying mechanism of the preferential and differential inclusion of the EDB exon during neoplastic transformation. Previous studies show that increased tissue stiffness directly upregulates PI3K/AKT-mediated SRp40 phosphorylation, enhancing exon inclusion and EDB-FN secretion by breast cancer cells (Bordeleau et al., 2015). In this work, development of drug resistance and TGF-β treatment in MCF7 and MDA-MB-468 cells consistently upregulated SRp55 phosphorylation, in addition to increased phospho-AKT and SRp40 levels. MK2206-HCl treatment showed highly specific knockdown of phospho-AKT and consequent downregulation of SRp40/SRp55 levels in a cell-specific manner. While the MK2206-HCl treatment significantly reduced EDB-FN expression and invasion, it did not completely abrogate them, suggesting the compensatory activation of other mitogenic proteins (like AKT3) (Stottrup et al., 2016) or the EDA-FN isoform, which is also involved in tumorigenesis (Han and Lu, 2017).

EDB-FN is overexpressed in multiple types of cancer, including breast, colorectal, oral, bladder, lung, and prostate (Bae et al., 2013, Lyons et al., 2001, Inufusa et al., 1995, Arnold et al., 2016, Khan et al., 2005, Albrecht et al., 1999). Originally thought to be secreted only by cancer-associated fibroblasts (CAFs) and endothelial cells, EDB-FN is now known to be abundantly produced by tumor cells, especially invasive tumor cells (Han and Lu, 2017). EDB-FN is upregulated during embryogenesis, temporally activated during wound healing, tissue repair, and angiogenesis, but mostly absent from healthy adult tissues (White et al., 2008). Additionally, by virtue of its extracellular location and ready accessibility, EDB-FN has emerged as an attractive target for designing new diagnostic and therapeutic regimens. Previous studies have already demonstrated the potential of antibody-mediated EDB-FN targeting (using L19, BC-1) for angiogenesis, inflammation, and cancer stem cell therapy (Mariani et al., 1997, Tijink et al., 2009). EDB-FN-specific peptides, such as ZD2 and APT_EDB_, are advantageous for oncogenic ECM targeting, by virtue of their small size, low immunogenicity, and high tissue penetration ability (Han et al., 2015, Sun et al., 2014, Zahnd et al., 2010). The specificity and superior binding of the ZD2 probe for EDB-FN has direct translational implications. We have successfully demonstrated differential diagnosis of non-invasive and invasive breast and prostate cancer xenografts in mouse models using ZD2-targeted MRI contrast agents (Han et al., 2017a, Han et al., 2018, Han et al., 2017b, Ayat et al., 2018). The results of this study open up new avenues for determining the potential of EDB-FN as a molecular biomarker for molecular imaging-based detection, risk-stratification, active surveillance, and monitoring of breast cancers and tracking their evolution as the disease progresses with and without chemotherapy.

In summary, this research demonstrates that EDB-FN expression is strongly associated with highly invasive breast cancer and with low-risk cells that evolve into high-risk cancer. Further, this correlation holds true despite cancer cell plasticity, and dynamic changes occurring in the invasive properties of breast cancer cells lead to corresponding changes in the EDB-FN expression levels. These observations indicate that EDB-FN is a promising molecular marker for monitoring the progression of breast cancer, in the context of diagnostic imaging and therapeutic interventions.

## Materials and Methods

### Cell lines and culture

MCF7, MDA-MB-231, BT549, and Hs578T cells were purchased from ATCC (Manassas, VA). MCF7-DR cells (resistant to 500 nM Palbociclib), MDA-MB-468, and MDA-MB-468-DR (resistant to 100 nM Paclitaxel) cells were a kind gift from Dr. Ruth Keri (CWRU, Cleveland, OH). MCF7-TGF-β and MDA-MB-468-TGF-β cells were obtained by treating the parent lines with 5 ng/mL TGF-β (RnD Systems, Minneapolis, MN) for at least 7-10 days. The breast cancer lines were cultured in Dulbecco’s Modified Eagle’s Medium (DMEM) supplemented with 10% fetal bovine serum (FBS), and 1% Penicillin/Streptomycin (P/S). MCF7, MCF7-DR, and Hs578T cells were additionally supplemented with 0.01 mg/mL human insulin (Sigma-Aldrich, St. Louis, MO). All the cells were grown at 37°C and 5% CO_2_. The cell lines were tested for the absence of mycoplasma using the MycoAlert™ Mycoplasma Detection Kit (Lonza, Allendale, NJ). Cell lines were also authenticated by Genetica DNA Laboratories (Burlington, NC).

### MK2206-HCl treatment

The invasive TGF-β-treated and drug-resistant MCF7 and MDA-MB-468 populations were treated with MK2206-HCl, a pan-AKT inhibitor (Agarwal et al., 2014), purchased from SelleckChem (Boston, MA). For this treatment, 8 × 10^5^ cells were plated on 6-well plates. After 24 h of attachment, the cells were treated with MK2206-HCl for 2 days (2 µM dose for MCF7 cells and 4 µM dose for MDA-MB-468 cells). Cells treated with equivalent volume of DMSO were used as controls. After treatment, a portion of the cells was harvested for protein and RNA extraction for western blotting and qRT-PCR. The rest of the cells were plated on Matrigel and in transwell inserts for the invasion and 3D growth assays, as described in the relevant sections, respectively.

### TCGA analysis

RNA-Seq data was mined from the TCGA database for the FN1 transcript containing the EDB-FN exon (ENST00000323926). Differential gene expression analysis was performed in 104 normal and 790 breast tumor samples. Statistical analysis and graph plotting was performed using Graphpad Prism.

### qRT-PCR

Total RNA was extracted from cells and tissues using the RNeasy Plus Mini Kit (Qiagen, Germantown, MD), according to manufacturer’s instructions. Reverse transcription was performed using the miScript II RT Kit (Qiagen) and qPCR was performed using the SyBr Green PCR Master Mix (Applied Biosystems, CA). Gene expression was analyzed by the 2^-ΔΔCt^ method with 18S and β-actin levels as the control. The following primer sequences were used, EDB-FN: Fwd 5’-CCGCTAAACTCTTCCACCATTA-3’ and Rev 5’-AGCCCTGTGACTGTGTAGTA-3’; FN1: Fwd 5’-CCTGGAGTACAATGTCAGTG-3’ and Rev 5’-GGTGGAGCCCAGGTGACA-3’; 18S: Fwd 5’-TCAAGAACGAAAGTCGGAGG-3’ and Rev 5’-GGACATCTAAGGGCATC ACA-3’; β-actin Fwd 5’-GTTGTCGACGACGAGCG-3’ and Rev 5’-AGCACAGAGCCTCGC CTTT-3’

### 3D tumor spheroid growth and ZD2-Cy5.5 staining

For the Matrigel growth assay, 5 × 10^5^ breast cancer cells were suspended in 5% Matrigel-containing media and plated on a thick layer of Corning™ Matrigel™ Membrane Matrix. The ability of the cells to form tumor spheroids in the 3D Matrix was monitored and photographed for up to 5 days using the Moticam T2 camera. After 5 days, the cells were stained with Hoechst (1:2000) and ZD2-Cy5.5 (100 nM) for 1 h. After 3 washes of PBS, fresh media was added and the cells were imaged by confocal microscopy on Olympus confocal microscope.

### Western blot

Total cellular protein was extracted as previously described (Vaidya et al., 2019). Protein extracts (40 µg) were run on SDS-PAGE, transferred onto nitrocellulose membrane and immunoblotted with primary antibodies overnight. The following primary antibodies (1:1000 dilution) were purchased from Cell Signaling Technology (Danvers, MA): anti-E-cadherin (Cat#3195), anti-Slug (Cat#9585), anti-phospho-T308-AKT (Cat#13038), anti-phospho-S473-AKT (Cat#4060), anti-pan-AKT (Cat#4691), and anti-β-actin (loading control, Cat#4970). The anti-Phosphoepitope SR proteins (Cat#MABE50; clone 1H4) and anti-SRp40 (Cat#06-1365) antibodies were purchased from Millipore Sigma (Temecula, CA) and used at 1:500 dilution. Anti-N-Cadherin antibody (Cat#76057) was purchased from Abcam (Cambridge, MA) and used at 1:500 dilution.

### Transwell assay

For the invasion assay, breast cancer cells were starved in serum-depleted media overnight. The next day, 1-2 × 10^5^ cells were plated in transwell inserts (VWR, Radnor, PA) coated with 0.3 mg/mL Corning™ Matrigel™ Membrane Matrix (Corning, NY). After 2 days, the inserts were swabbed with Q-tips to remove the plated cells. The invading cells on the bottom of the inserts were fixed with 4% paraformaldehyde followed by staining with 0.1% crystal violet for 20 min. Excess stain was washed under tap water and images of the purple migrated cells were taken using the Moticam T2 camera.

### EDB-FN knockdown

ECO/siRNA nanoparticles were formulated as previously described (Gujrati et al., 2016). Briefly, the cationic lipid ECO (5 mM stock in ethanol) was mixed with siEDB or siLuc (as negative control NC) at a final siRNA concentration of 100 nM and N/P = 10 for 30 min to enable self-assembly formation of ECO/siEDB or ECO/NC nanoparticles, respectively. For transfections, the nanoparticle formulation was mixed with culture media and added on to MDA-MB-468-DR cells. After 48 h, the cells were stained with ZD2-Cy5.5 as described above. The siEDB duplex [sense 5’-GCA UCG GCC UGA GGU GGA CTT-3’ and antisense 5’-GUC CAC CUC AGG CCG AUG CTT-3’] was purchased from IDT (Coralville, IA). The siLuc duplex [sense 5’-CCU ACG CCG AGU ACU UCG AdTdT-3’ and antisense 5’-dTdT GGA UGC GGC UCA UGA AGC U-3’] was purchased from Dharmacon (Lafayette, CO).

### Statistical analyses

All the experiments were independently performed in triplicates (n = 3). Data are represented as mean ± s.e.m. Statistical analysis was performed using Graphpad Prism. Data between two groups was compared using unpaired Student’s *t*-test. p < 0.05 was considered to be statistically significant.

## Abbreviations

BCa: Breast cancer
EDB-FN: Extradomain-B fibronectin
FN1: Fibronectin
MRI: Magnetic resonance imaging
PET: Positron-emission tomography
CT: Computed tomography
TME: Tumor microenvironment
ECM: Extracellular matrix
TGF-β: Transforming growth factor-β
EMT: Epithelial to mesenchymal transition
CEA: Carcinoembryonic antigen

## Acknowledgements

We are thankful to Dr. Zheng Han for providing the ZD2-Cy5.5 probe and to Dr. Nadia Ayat for her helpful comments.

## Contributions

The conceptual design of the research was devised by AMV and ZRL. Experimental execution of all aspects of the research was done by AMV. VQ and HW participated in cell culture, qRT-PCR, western blot, and *in vitro* functional assays. The manuscript was written and edited by AMV and ZRL.

